# Deubiquitinase USP15 restricts autophagy and macrophage immunity to *Mycobacterium tuberculosis*

**DOI:** 10.1101/2025.08.06.668987

**Authors:** Kathryn C. Rahlwes, Priscila C. Campos, Beatriz R. S. Dias, Paola K. Párraga, Michael U. Shiloh

**Affiliations:** Department of Internal Medicine, University of Texas Southwestern Medical Center, Dallas, Texas, USA; Department of Microbiology, University of Texas Southwestern Medical Center, Dallas, Texas, USA

## Abstract

Autophagy enables macrophages to degrade intracellular *Mycobacterium tuberculosis* (Mtb), and this defense depends on E3 ubiquitin ligases such as PARKIN and SMURF1, which tag Mtb-associated structures for lysosomal clearance. Deubiquitinases (DUBs) counter ubiquitin ligases by removing ubiquitin from molecular targets. We hypothesized that DUBs might offset ubiquitin ligase activity and negatively regulate host immunity to Mtb. Here, we identify ubiquitin-specific protease 15 (USP15) as a negative regulator of autophagy-mediated macrophage immunity to Mtb. Using a targeted knockdown screen in mouse macrophages, we found that *Usp15* loss increased K63-linked ubiquitination and LC3 recruitment to Mtb-associated structures, leading to reduced bacterial replication. These effects required USP15’s catalytic activity and were reversed by knockdown of PARKIN (*Park2*) or inhibition of autophagy initiation. In primary human macrophages, *USP15* knockdown similarly enhanced LC3 targeting and restricted Mtb growth. Importantly, pharmacologic inhibition of USP15 with a selective small molecule decreased Mtb burden in human macrophages. Our findings identify USP15 as a suppressor of macrophage immunity and suggest that targeting deubiquitinases may represent a promising host-directed therapeutic strategy against tuberculosis.

## Introduction

*Mycobacterium tuberculosis* (Mtb), the causative agent of tuberculosis (TB), is responsible for an estimated 10 million new cases and 1.3 million deaths annually worldwide (1). The airborne route of transmission, combined with its ability to persist within infected individuals, contribute to its global success (2, 3). Mtb primarily infects macrophages or inflammatory monocytes following inhalation (4, 5) and can enter a long-lived persistent state that later reactivates to cause disease and transmission (3, 6). While antibiotics are effective against active TB, they are less effective against latent bacteria, and the emergence of multidrug-resistant Mtb strains further complicates treatment (1, 7). These challenges underscore the need for host-directed therapies that enhance immune control of Mtb replication (8–14). However, a detailed understanding of the molecular mechanisms used by host innate immune cells to restrict bacterial replication is incomplete.

Xenophagy, a form of selective autophagy that targets large intracellular structures, is a critical component of macrophage cell-autonomous immunity against Mtb (15–18). Pathogen- and damage-associated molecular patterns (PAMPs and DAMPs) stimulate xenophagy by recruiting E3 ubiquitin ligases such as PARKIN (encoded by the *Park2* gene) and SMURF1 to cytosolic Mtb or damaged phagosomal membranes (19–26). These ligases attach ubiquitin to Mtb-associated structures, particularly K63- and K48-linked polyubiquitin chains, which are recognized by adaptor proteins like p62, NDP52, NBR1 and Optineurin (OPTN). In turn, adaptors recruit LC3 and other autophagy machinery, leading to the formation of autophagosomes and subsequent bacterial degradation through lysosomal fusion (27–29). While some controversy remains regarding the role of autophagy in Mtb control (30, 31), genetic ablation of *Park2*, *Smurf1*, or autophagy pathway components in murine (25, 26, 32–35) or human (26, 36) macrophages results in increased Mtb growth and reduced mouse survival (25, 26, 32–35).

Although the role of ubiquitin ligases in promoting xenophagy is well established, the contribution of opposing deubiquitinases (DUBs) remains poorly understood. DUBs remove ubiquitin chains from proteins and are categorized into five major families based on structural and functional characteristics (37). These enzymes are known to regulate key immune and stress response pathways, but only a few have been implicated in macrophage immunity during infection. For example, DUB30 negatively regulates mitophagy by delaying mitochondrial recruitment of PARKIN (38, 39), and ubiquitin-specific protease (USP) 18 modulates type I interferon signaling and host defense during Mtb and *Salmonella* infections (40). Finally, a recent study identified USP8 as a negative regulator of cell-autonomous immunity to Mtb by facilitating the repair of Mtb-induced membrane damage that would normally trigger Mtb degradation (41). However, the broader roles of DUBs in regulating host responses to Mtb remain largely unexplored.

To address this gap, we performed a knockdown screen targeting murine DUBs in BV2 macrophages (42) and identified multiple candidates that alter Mtb replication. USP15 emerged as a potent negative regulator of host immunity to Mtb. Genetic deletion or knockdown of USP15 in mouse and human macrophages enhanced K63-linked ubiquitination and LC3 recruitment to Mtb-associated structures, resulting in impaired bacterial replication. These effects required USP15’s catalytic activity and were reversed by knockdown of *Park2* or pharmacologic inhibition of autophagy initiation. In primary human macrophages, pharmacologic inhibition of USP15 similarly reduced Mtb burden. These findings identify USP15 as a key regulator of autophagy and macrophage defense against Mtb and suggest that targeting deubiquitinases may represent a promising strategy for host-directed therapy against tuberculosis.

## Results

### Depletion or deletion of *Usp15* inhibits Mtb replication

To identify deubiquitinases (DUBs) that regulate Mycobacterium tuberculosis (Mtb) replication in macrophages, we generated a knockdown library in the genetically tractable murine BV2 macrophage cell line. We selected BV2 microglial macrophages due to their robust genetic tractability, their previously validated use in autophagy studies, and their established utility in modeling mycobacterial pathogenesis (43–50). We used a luminescent Mtb strain (Mtb-pLux) to measure intracellular bacterial growth across cell lines, each stably expressing one of three distinct shRNAs per target gene (Figure 1A). Among the DUBs screened, knockdown of 33 genes increased Mtb replication, whereas knockdown of 6 genes decreased bacterial growth by at least 1.25-fold compared to a non-targeting control (NTC) (Figure 1A, 1B). Notably, *Usp18* knockdown increased Mtb growth, consistent with prior reports and validating our screening strategy (40).

**Figure 1.**
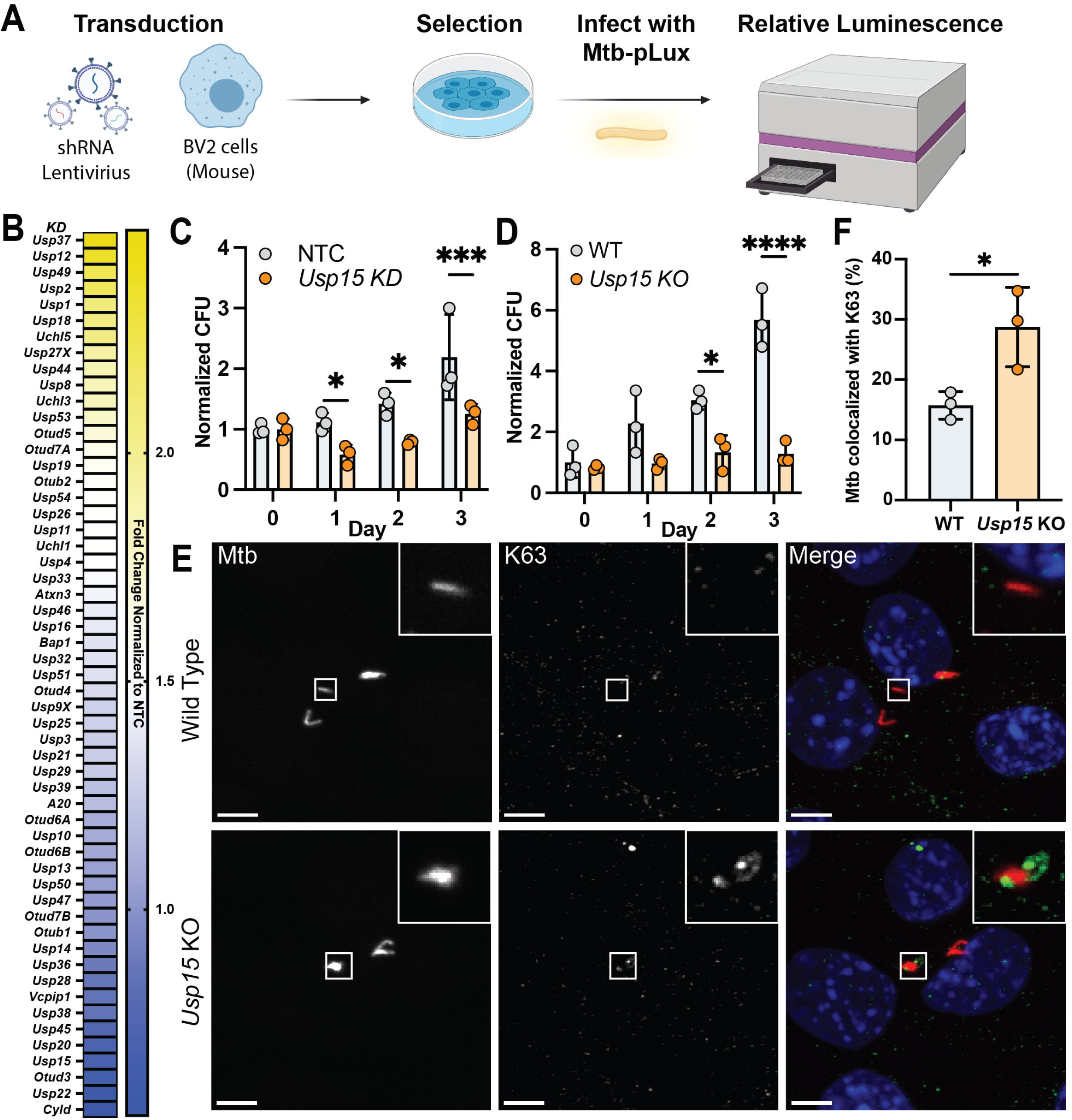
Loss of *Usp15* inhibits Mtb replication and increases K63 ubiquitination co-localization. A) Schematic for shRNA screen in the BV2 cell line infected with luminescent Mtb. B) Heat map shows the average of relative luminescence units when normalized to day 0 and then to the nontargeting control (NTC). C) Colony forming unit (CFU) values of WT Erdman Mtb infected into non-targeting control shRNA (NTC) and *Usp15* KD shRNA BV2 cells were normalized for each day to the count on Day 0. D) CFUs of WT or CRISPR *Usp15* KO BV2 infected with WT Mtb. E) Images show Mtb (gray or red) and K63 (gray or green) with DAPI (blue) in WT or *Usp15* KO BV2 cells. Scale bar is 5 µM. F) Quantification of K63 immunofluorescence co-localizing with mCherry Mtb infected WT or *Usp15* KO BV2 cells at 16h post-infection. For statistical analysis of CFU, we used two-way ANOVA and Tukey’s multiple comparison test. For statistical analysis of co-localization, we used Student’s t-test. * p<0.05, ** p<0.01, *** p<0.001, **** p<0.0001. Data shown are representative of at least 3 independent experimental replicates.

We focused on USP15 based on its significant suppression of Mtb replication and its known interactions with PARKIN and roles in autophagy regulation (51–53). We first validated *Usp15* knockdown (KD) in BV2 macrophages using two independent shRNAs, which effectively reduced USP15 protein levels by immunoblotting (Supplemental Figure 1A). We next quantified Mtb replication in BV2 *Usp15* KD cell lines using a traditional colony-forming unit (CFU) assay. Compared to control BV2 cells, BV2 *Usp15* KD macrophages exhibited static Mtb growth and significantly reduced CFU over a 3-day infection (Figure 1C).

To confirm this phenotype, we generated BV2 *Usp15* knockout (KO) cells using CRISPR/Cas9 and validated USP15 loss by sequencing and immunoblotting (Supplemental Figure 1B). In agreement with the knockdown data, BV2 *Usp15* KO macrophages infected with Mtb Erdman (MOI 1) showed reduced CFU at two and three days post-infection compared to wild-type BV2 macrophages (Figure 1D). Bacterial burden remained static in BV2 *Usp15* KO cells over time, while it increased steadily in wild-type controls. These results demonstrate that *Usp15* suppresses cell-autonomous immunity to Mtb in BV2 macrophages.

### *Usp15* deletion in BV2 cells increases K63 ubiquitin co-localization with Mtb

Because USP15 has been reported to remove K63-linked ubiquitin (K63-Ub) and counteract PARKIN activity (51), we next tested whether loss of *Usp15* affected K63 ubiquitination of Mtb-associated structures. We use the term “Mtb-associated structures” because it remains unclear whether ubiquitin directly attaches to bacteria or the membranes surrounding the bacteria. We infected wild-type and BV2 *Usp15* KO cells with mCherry-expressing Mtb and performed immunofluorescence microscopy using an anti-K63 ubiquitin antibody. Compared to wild-type cells, BV2 *Usp15* KO cells exhibited a significant increase in the colocalization of K63-Ub with Mtb (Figure 1E, 1F). As a control, we examined K48-linked ubiquitin (K48-Ub), which has also been reported to localize to Mtb-associated structures (25). In contrast to K63-Ub, K48-Ub colocalization with mCherry Mtb was unchanged between wild-type and BV2 *Usp15* KO cells (Supplemental Figure 1C, 1D). These results suggest that USP15 specifically limits the accumulation of K63-linked ubiquitin at Mtb-associated structures, consistent with its role in countering PARKIN-mediated ubiquitination.

### *Usp15* deletion in BV2 cells increases LC3-I to LC3-II conversion and recruitment of LC3 to Mtb-associated structures

Because K63-linked ubiquitination has been implicated in promoting recruitment of autophagy machinery (25), we next tested whether loss of *Usp15* altered LC3 dynamics during Mtb infection. One hallmark of autophagy activation is the conversion of cytosolic LC3-I to the lipidated LC3-II form, which becomes membrane-associated during autophagosome formation (54, 55). BV2 *Usp15* KO cells infected with Mtb exhibited increased LC3-II relative to LC3-I, as measured by immunoblotting (Figure 2A, 2B). We next examined whether this increase in LC3-II was accompanied by enhanced recruitment of LC3 to Mtb-associated structures. Immunofluorescence microscopy revealed greater colocalization of endogenous LC3 with mCherry Mtb in BV2 *Usp15* KO cells compared to wild-type controls (Figure 2C, 2D).

**Figure 2.**
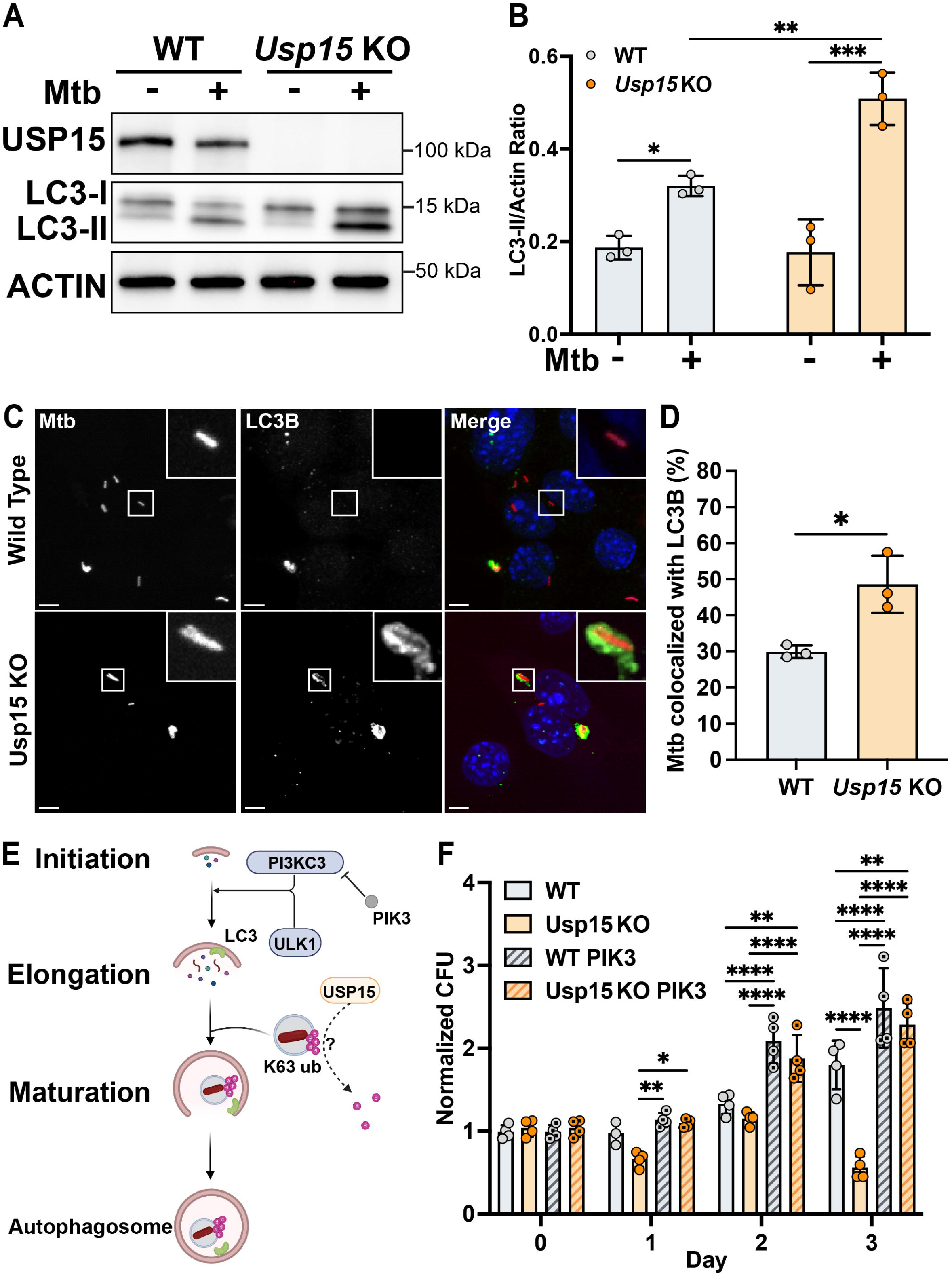
*Usp15* deletion in BV2 cells increases LC3-II conversion and colocalization of LC3 with Mtb-associated structures. A) Western blot of LC3-I and LC3-II and ACTIN in WT and *Usp15* KO BV2 cells infected with Mtb. B) Quantification of LC3-II normalized to ACTIN. C) Representative immunofluorescence images of mCherry Mtb (grey or red), LC3 (grey or green) in WT (upper panel) or *Usp15* KO (lower panel) BV2 cells. Scale bar is 5 µM. D) Quantification of LC3 co-localization with mCherry Mtb in BV2 cells. E) Schematic of PIK-III inhibition of upstream autophagy initiation via the PI3KC3 complex. F) CFU of WT Mtb in WT or *Usp15* KO cells with or without 5 µM of PIK-III normalized for each day to the count on Day 0. For statistical analysis, we used two-way ANOVA and Tukey’s multiple comparison test.* p<0.05, ** p<0.01, *** p<0.001, **** p<0.000.1. Data shown are representative of at least 3 independent experimental replicates.

To determine whether this phenotype was dependent on canonical autophagy initiation, we treated wild-type and BV2 *Usp15* KO macrophages with PIK-III, an inhibitor of the phosphoinositide 3-kinase complex component VPS34 essential for autophagy initiation (Figure 2E) (56). As expected, inhibition of autophagy increased Mtb replication in wild-type BV2 cells (Figure 2F). Importantly, PIK-III treatment also reversed the reduced Mtb burden in BV2 *Usp15* KO cells (Figure 2F), indicating that autophagy is required for the enhanced bacterial control observed in the absence of *Usp15* in macrophages.

### *Usp15* deletion in mouse bone marrow-derived macrophages leads to decreased bacterial replication and increased autophagy

To determine whether *Usp15* deletion affects macrophage immunity in primary cells, we examined bone marrow-derived macrophages (BMDMs) from wild-type and *Usp15* knockout mice (57, 58). Following infection with Mtb, BMDMs from *Usp15* KO mice exhibited significantly reduced bacterial burden compared to wild-type controls at day 3 post-infection (Figure 3A). We next tested whether *Usp15* deletion in BMDMs enhanced autophagy activation. Immunofluorescence microscopy showed that endogenous LC3 was more frequently colocalized with mCherry Mtb in *Usp15* KO BMDMs (Figure 3B, 3C), consistent with enhanced recruitment of autophagy machinery. Together, these results confirm that the autophagy-dependent restriction of Mtb observed in BV2 cells also occurs in primary mouse macrophages lacking *Usp15*.

**Figure 3.**
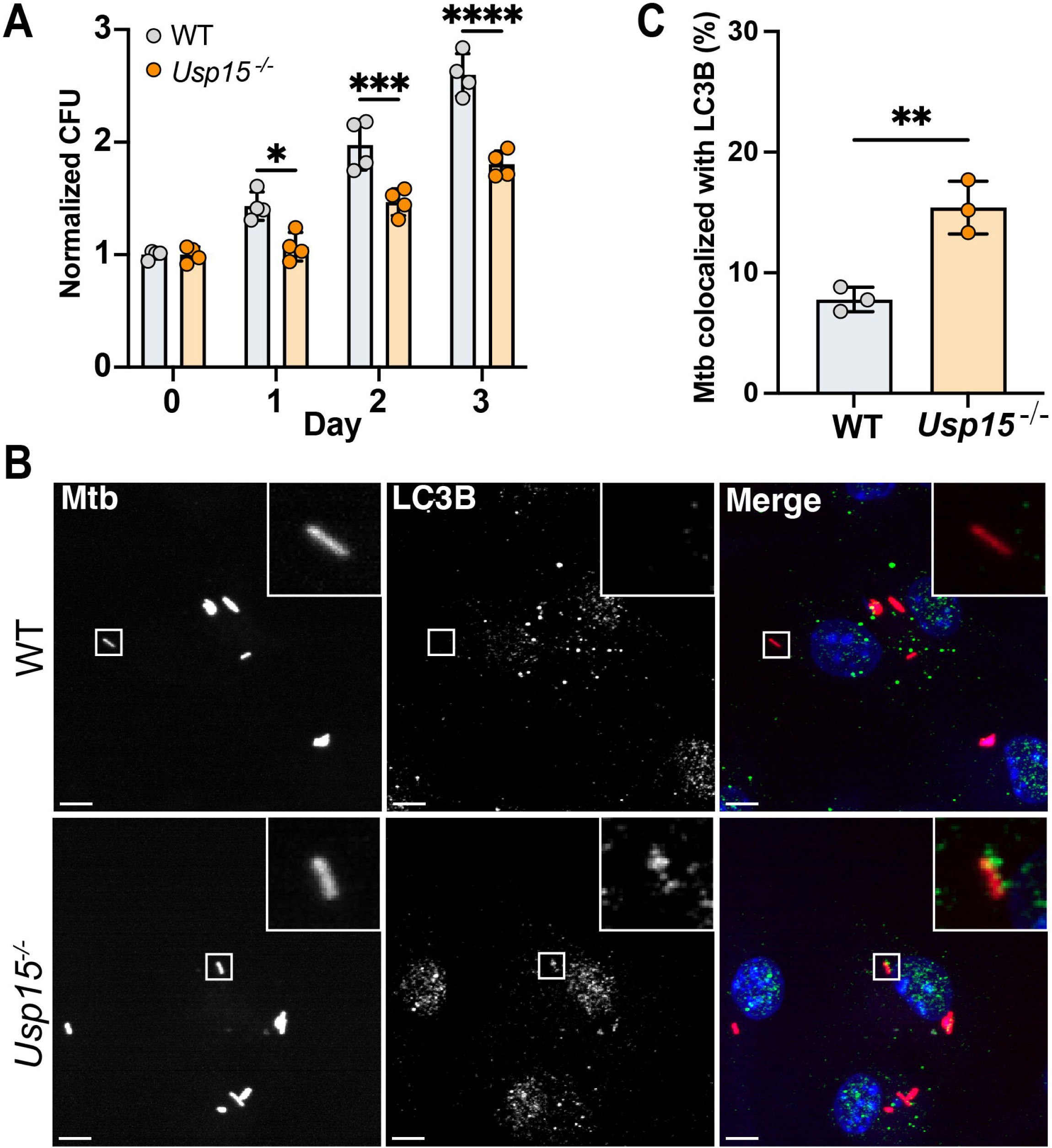
Loss of *Usp15* in BMDM results in reduced CFU and increased co-localization of LC3 with Mtb-associated structures. A) CFU of WT Mtb in WT or *Usp15*^−/-^ BMDMs normalized to the count at Day 0. B) Representative images of mCherry (grey or red) and LC3 immunofluorescence (grey or green) in WT or *Usp15*^−/-^ BMDMs. Scale bar is 5 µM. C) Quantification of LC3 co-localization with mCherry Mtb in BMDMs 16 at 16h post-infection. For statistical analysis of CFU, we used two-way ANOVA and Tukey’s multiple comparison test. For statistical analysis of co-localization, we used Student’s t-test. * p<0.05, ** p<0.01, *** p<0.001, **** p<0.0001. Data shown are representative of at least 3 independent experimental replicates.

### The deubiquitinase activity of USP15 is necessary for its role in regulating Mtb replication in BV2 macrophages

USP15 contains a conserved catalytic triad (Cys269, His862, Asp879) typical of the USP family of deubiquitinases (59). To determine whether USP15 suppresses macrophage immunity to Mtb through its enzymatic function, we generated BV2 *Usp15* KO macrophages stably complemented with either wild-type (WT) USP15 or a catalytically inactive mutant, C269A, which targets the catalytic cysteine within the USP domain triad (59). Specifically, we complemented BV2 *Usp15* KO cells with either an empty vector, a vector containing a cDNA encoding full-length USP15 (*Usp15^WT^*), or a vector containing a cDNA encoding the catalytically inactive USP15^C269A^ allele, each also containing an N-terminal 3X-FLAG tag (Figure 4A, 4B). Complementation with WT USP15 fully restored intracellular Mtb replication to levels observed in parental BV2 cells, whereas expression of the USP15^C269A^ mutant did not (Figure 4C).

**Figure 4.**
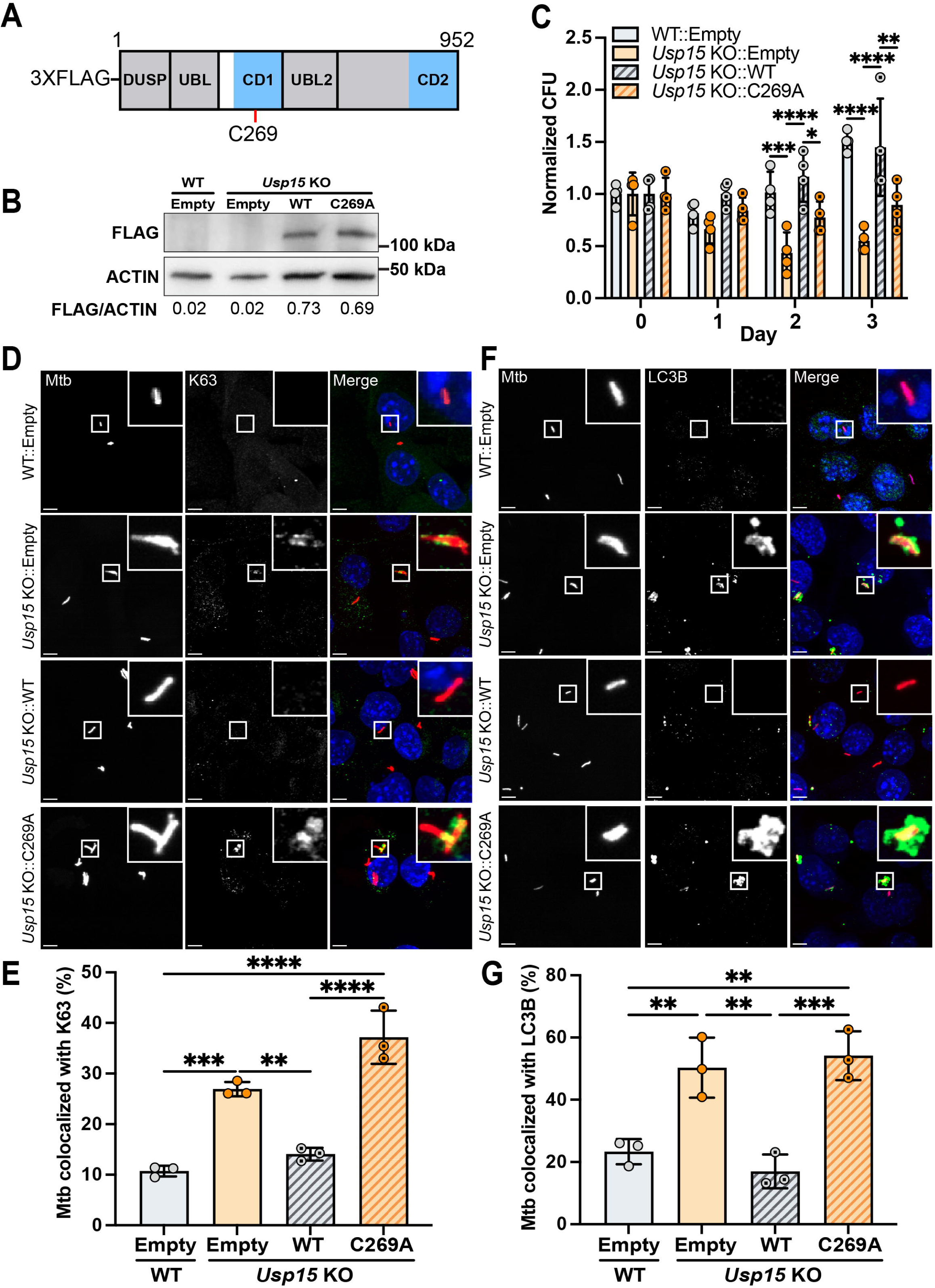
The catalytic activity of *USP15* is necessary for regulating Mtb replication in BV2 cells. A) Schematic of *Usp15* with the location of the catalytic dead mutation noted in red. Domain abbreviations are as follows: DUSP (domain present in USP) and UBL (ubiquitin-like domain). The catalytic domains (CD) are noted in blue. B) Complementation of BV2 *Usp15* KO cells with 3XFlag-Usp15WT (Usp15 KO::WT) or 3XFlag-Usp15C269A (*Usp15* KO::C269A). Western blot demonstrating 3XFlag-*Usp15*^WT^ or 3XFlag -*Usp15*^C269A^ in BV2 cell lines. C) CFU of Mtb infection in BV2 WT with empty vector (WT Empty), BV2 *Usp15* KO with empty vector (*Usp15* KO Empty), BV2 *Usp15* KO complemented with 3XFlag-*Usp15*^WT^ (*Usp15* KO::WT) or BV2 *Usp15* KO complemented with 3XFlag-*Usp15*^C269A^ (*Usp15* KO:: C269A). CFU values were normalized to Day 0. D) Representative image of mCherry (grey or red) and K63 immunofluorescence (grey or green) with DAPI (blue) in WT Empty, BV2 *Usp15* KO Empty, BV2 *Usp15* KO::WT, or BV2 *Usp15* KO::C269A. Scale bar is 5 µM. E) Quantification of K63 co-localization with mCherry Mtb in WT Empty, BV2 *Usp15* KO Empty, BV2 *Usp15* KO::WT, or BV2 *Usp15* KO::C269A. F) Representative image of mCherry (grey or red) and LC3 immunofluorescence (grey or green) with DAPI (blue) in WT Empty, BV2 *Usp15* KO Empty, BV2 *Usp15* KO::WT, or BV2 *Usp15* KO::C269A. Scale bar is 5 µM. G) Quantification of LC3 co-localization with mCherry Mtb in WT Empty, BV2 *Usp15* KO Empty, BV2 *Usp15* KO::WT, or BV2 *Usp15* KO::C269A. For statistical analysis of CFU, we used two-way ANOVA and Tukey’s multiple comparison test. For statistical analysis of colocalization, we used one-way ANOVA and Tukey’s multiple comparison test. * p<0.05, ** p<0.01, *** p<0.001, **** p<0.0001. Data shown are representative of at least 3 independent experimental replicates.

We next examined whether USP15 enzymatic activity was required to suppress K63-ubiquitination and LC3 recruitment to Mtb-associated structures. Immunofluorescence microscopy showed that only WT USP15 reversed the increased K63-ubiquitination and LC3 colocalization with mCherry Mtb observed in BV2 *Usp15* KO cells, while the USP15^C269A^ mutant failed to do so (Figure 4D-4G). These results indicate that USP15 requires its catalytic activity to restrict autophagy and promote intracellular Mtb growth in macrophages.

### USP15 counters the activity of PARKIN on K63 ubiquitination and Mtb survival

Previous studies suggest that USP15 interacts with and counteracts PARKIN, an E3 ubiquitin ligase that promotes K63-linked polyubiquitination of Mtb and enhances autophagy-mediated clearance (Figure 5A) (25, 26, 51). We therefore tested whether the effects of *Usp15* deletion on Mtb ubiquitination and intracellular growth were dependent on PARKIN. First, we performed shRNA-mediated knockdown of *Park2* in BV2 *Usp15* KO cells and confirmed loss of PARKIN expression by immunoblotting (Figure 5B). As expected, *Park2* knockdown in wild type BV2 cells resulted in increased bacterial burden over time compared to wild type BV2 cells expressing the non-targeting control (Figure 5C). In addition, compared to non-targeting controls, *Park2* knockdown restored Mtb replication in BV2 *Usp15* KO cells nearly to levels observed in wild-type BV2 cells at 3 days after infection (Figure 5C).

**Figure 5.**
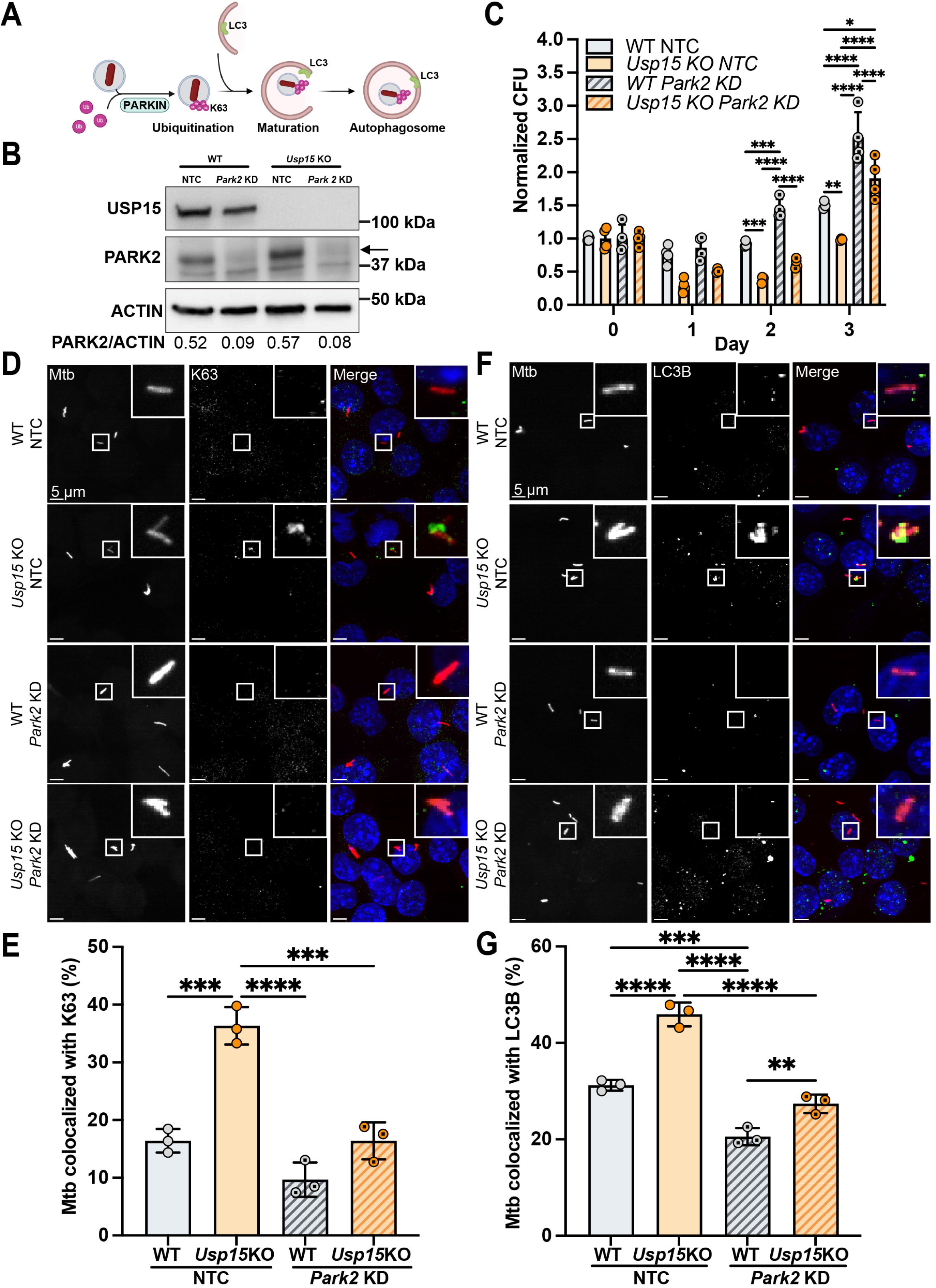
USP15 counters the activity of PARKIN. A) Schematic of PARKIN’s activity in the context of ubiquitination of Mtb. B) Western blot confirms knockdown (KD) of PARKIN in WT or *Usp15* KO compared to non-targeting controls (NTC). The percentage of KD is shown below each KD. C) CFU of WT Mtb infected in WT with NTC, *Usp15* KO with NTC, WT with *Parkin* KD, and *Usp15* KO with *Parkin* KD. CFU values were normalized to the count at Day 0. D) Representative image of mCherry (grey or red) and K63 immunofluorescence (grey or green) with DAPI (blue) in WT with NTC, *Usp15* KO with NTC, WT with *Parkin* KD, and *Usp15* KO with *Parkin* KD. Scale bar is 5 µM E) Quantification of K63 co-localization with mCherry Mtb in WT with NTC, *Usp15* KO with NTC, WT with *Parkin* KD, and *Usp15* KO with *Parkin* KD. F) Representative image of mCherry (grey or red) and LC3 immunofluorescence (grey or green) with DAPI (blue) in WT with NTC, *Usp15* KO with NTC, WT with *Parkin* KD, and *Usp15* KO with *Parkin* KD. Scale bar is 5 µM. G) Quantification of LC3 co-localization with mCherry Mtb in WT with NTC, *Usp15* KO with NTC, WT with *Parkin* KD, and *Usp15* KO with *Parkin* KD. For statistical analysis of CFU, we used two-way ANOVA and Tukey’s multiple comparison test. For statistical analysis of colocalization, we used one-way ANOVA and Tukey’s multiple comparison test. * p<0.05, ** p<0.01, *** p<0.001, **** p<0.0001. Data shown are representative of at least 3 independent experimental replicates.

Next, we examined the impact of *Park2* knockdown on K63-linked ubiquitination and LC3 recruitment to Mtb associated structures. Immunofluorescence microscopy showed that in BV2 *Usp15* KO cells expressing a *Park2* shRNA, the frequency of K63-ubiquitin-positive and LC3-positive Mtb-associated structures was significantly reduced compared to BV2 *Usp15* KO cells encoding a non-targeting control (Figure 5D-5G). These findings demonstrate that USP15 restricts macrophage immunity to Mtb, at least in part, by opposing PARKIN-mediated K63 ubiquitination and downstream engagement of autophagy machinery components.

### *USP15* depletion in human monocyte-derived macrophages leads to decreased Mtb replication

To further validate the role of USP15 in cell-autonomous immunity to Mtb, we tested the impact of genetic depletion of *USP15* on Mtb replication in primary human monocyte-derived macrophages (hMDM). We isolated peripheral blood mononuclear cells (PBMCs) from buffy coat preparations obtained from healthy blood donors, knocked down *USP15* through lentiviral transduction of *USP15*-specific shRNA, and differentiated the cells into macrophages over seven days. We achieved approximately 40–50% knockdown of USP15 as determined by immunoblot (Figure 6A), which is consistent with our prior results targeting *SMURF1* (26).

**Figure 6.**
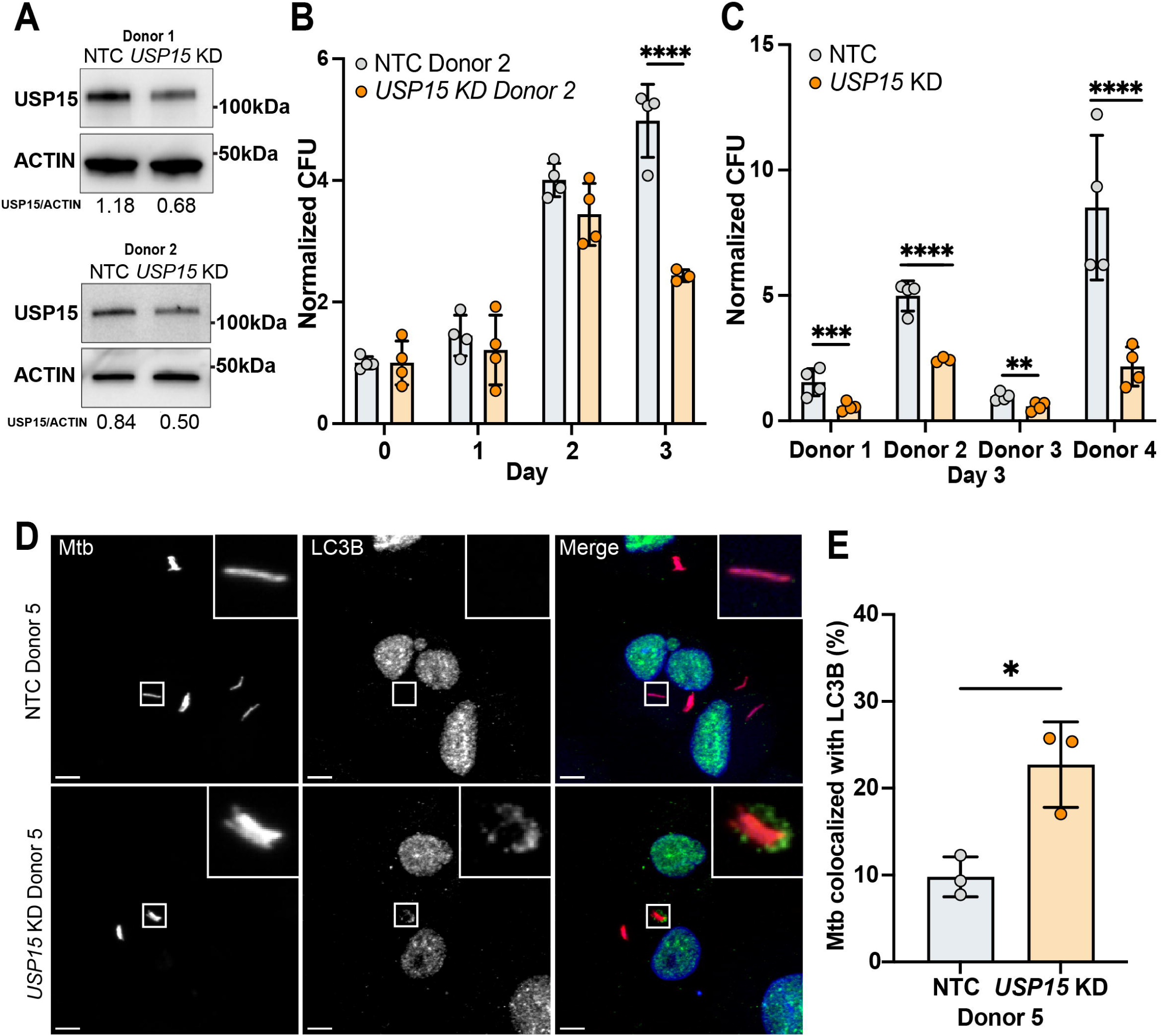
USP15 depletion in hMDM leads to decreased Mtb burden and increased LC3 co-localization. A) Representative western blot of knockdown of *USP15* in human monocyte-derived macrophages (hMDMs) from Donor 1 and Donor 2. B) Representative time course from day 0 to day 3 of Mtb CFU in hMDMs from Donor 2. C) Combined normalized CFU to day 0 from day 3 of Donor 1 (D1), Donor 2 (D2), Donor 3 (D3), and Donor 4 (D4). D) Representative image of mCherry Mtb (grey or red) and LC3 immunofluorescence (grey or green) in hMDMs from Donor 5 (D5) with NTC or *USP15* KD. Scale bar is 5 µM. E) Quantification of LC3 co-localization in hMDMs from Donor 5 with NTC or *USP15* KD. For statistical analysis of CFU, we used two-way ANOVA and Tukey’s multiple comparison test. For statistical analysis of colocalization, we used Student’s t-test.* p<0.05, ** p<0.01, *** p<0.001, **** p<0.0001. Data shown are representative of at least 3 independent experimental replicates.

Despite partial knockdown, hMDMs from multiple donors transduced with *USP15*-specific shRNA exhibited reduced Mtb CFU compared to those transduced with a non-targeting control shRNA (Figure 6B, 6C). To determine whether this phenotype correlated with autophagy induction, we assessed LC3 colocalization with Mtb-associated structures by immunofluorescence. *USP15* knockdown resulted in increased LC3 recruitment relative to control cells (Figure 6D, 6E; Supplemental Figure 2). We also attempted to assess K63-linked ubiquitination of Mtb-associated structures in hMDMs using multiple antibodies but were unable to detect K63-Ub reliably by immunofluorescence, preventing us from evaluating whether *USP15* knockdown altered ubiquitination dynamics in this setting.

Together, these data show that USP15 suppresses autophagy-mediated control of Mtb in primary human macrophages, further supporting its role as a negative regulator of macrophage cell-autonomous immunity in both murine and human systems.

### Pharmacologic inhibition of USP15 leads to decreased bacterial replication

Based on the findings that USP15 suppresses antibacterial autophagy in both mouse and human macrophages, we next tested whether pharmacologic inhibition of USP15 could recapitulate these effects. We used a recently described small molecule inhibitor, USP15-IN-1 (60), to assess whether pharmacologic inhibition of USP15 restricts Mtb replication in macrophages. First, we confirmed that USP15-IN-1 had no direct effect on axenic Mtb growth (Supplemental Figure 3A). Next, to test specificity of USP15-IN-1, we treated wild-type and BV2 *Usp15* KO cells with increasing concentrations of USP15-IN-1 and infected them with Mtb-pLux. In wild-type BV2 cells, USP15-IN-1 reduced Mtb replication by approximately two-fold at the highest concentration tested (60 µM) (Figure 7A). This reduction was not observed in BV2 *Usp15* KO cells, indicating the effect was USP15-dependent. Cell viability was unaffected by USP15-IN-1 at all tested concentrations (Supplemental Figure 3B, 3C).

**Figure 7.**
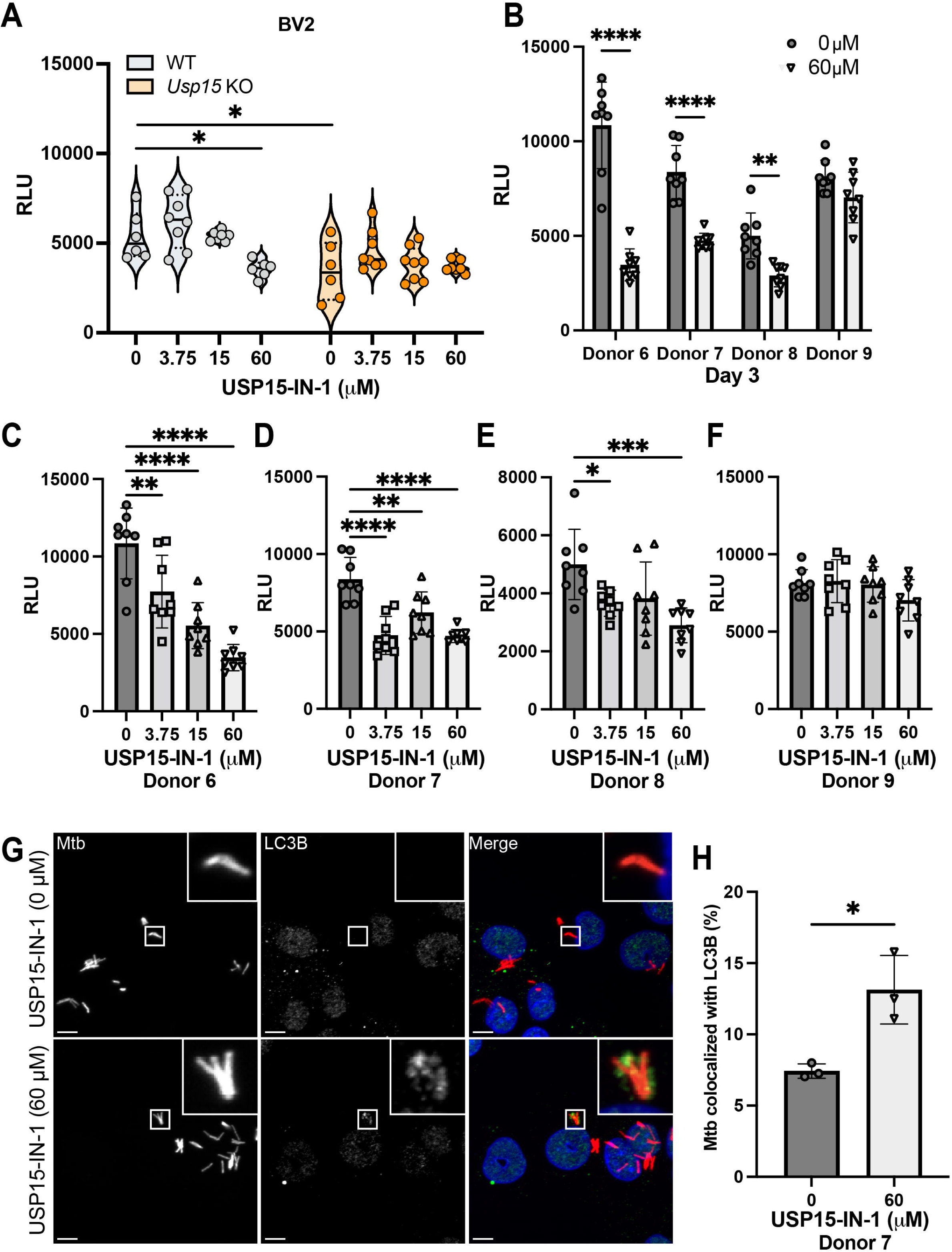
Inhibition of USP15 by USP15-IN-1 leads to decreased Mtb burden in BV2 cells and hMDM. A) A 4-point dose response of USP15-IN-1 in WT or Usp15 KO BV2 cells infected with Mtb-pLux. B) Combined Relative Luminescence Units (RLU) of Day 3 from Donor 6, Donor 7, Donor 8, and Donor 9. C-F) The 4-point dose response of USP15-IN-1 in C) Donor 6, D) Donor 7, E) Donor 8, F) Donor 9. G) Representative immunofluorescence image of mCherry Mtb (grey or red) and LC3 (grey or green) in hMDMs from Donor 7 with 0 µM (DMSO control) or 60 µM of 15 IN 1 at 18 hours post-infection. Scale bar is 5 µM. H) Quantification of LC3 co-localization in hMDMs from donor 7. For statistical analysis for Mtb-pLux experiments, we used two-way ANOVA and Tukey’s multiple comparison test. For colocalization analysis, we used Student’s t-test. * p<0.05, ** p<0.01, *** p<0.001, **** p<0.0001. Data shown are representative of at least 3 independent experimental replicates.

We next tested the efficacy of USP15-IN-1 in hMDMs derived from four healthy donors. USP15-IN-1 treatment resulted in a dose-dependent decrease in Mtb replication in three of four donors (Figure 7B-F). Notably, Mtb replication was reduced by nearly 50% with as little as 3.75 µM USP15-IN-1 in responsive donors (Figure 7C-E), without evidence of cytotoxicity as determined by LDH release assay (Supplemental Figure 3D, 3E).

To assess whether these effects were associated with autophagy induction, we evaluated LC3 colocalization with Mtb. USP15-IN-1 increased LC3 recruitment in hMDMs from two of the three donors (Figure 7G, 7H; Supplemental Figure 3F-I). No increase in LC3 recruitment was observed in hMDMs from donor 9, which also failed to respond to USP15-IN-1 with reduced bacterial replication (Figure 7F; Supplemental Figure 3F, 3G).

Together, these data demonstrate that pharmacologic inhibition of USP15 with USP15-IN-1 enhances autophagy-mediated restriction of Mtb in human macrophages, supporting its potential as a host-directed therapeutic strategy for tuberculosis.

## Discussion

Host-directed therapies that harness or enhance innate immune pathways offer a promising avenue to improve outcomes in tuberculosis (TB), particularly in the face of rising drug resistance (61–63). In this study, we identify the deubiquitinase USP15 as a conserved and targetable suppressor of autophagy-mediated immunity to Mtb in both murine and human macrophages. Our findings reveal that USP15 removes PARKIN-dependent K63-linked ubiquitin from Mtb-associated structures, reducing LC3 recruitment and autophagic clearance. Genetic deletion or pharmacologic inhibition of USP15 increases K63 ubiquitination, enhances LC3 colocalization, and restricts Mtb replication in multiple macrophage systems. While USP15 has been linked to mitophagy and mitochondrial quality control through interactions with PARKIN (38, 51), our study newly defines its role in pathogen-directed autophagy, extending current understanding of USP15’s role in host-pathogen interactions.

Beyond its role in xenophagy, USP15 may regulate additional immune signaling pathways relevant to Mtb infection (64, 65). For instance, USP15, with the aid of COP9 signalosome (CSN), is a negative regulator of NF-κB via deubiquitination of IκBα (66, 67). Furthermore, loss of USP15 has been suggested to impact lipid droplet accumulation while decreasing HCV virus propagation (52). Lipid droplets are considered an additional nutrient source that can influence the balance of Mtb between dormancy and active growth (68, 69). In addition, USP15 stabilizes TRIM25 to enhance production of IFN-β (70, 71), which itself is considered a key regulator of Mtb replication (72). Thus, USP15 may act as a direct negative regulator of xenophagy and influence additional signaling pathways during Mtb infection.

By comparing our screen to that of Chandra *et al*. (41), we validate USP15 as a key negative regulator of macrophage immunity and highlight the broader relevance of deubiquitinases in host-pathogen interactions. Notably, Chandra *et al.* showed that knockdown of *Usp15* in immortalized BMDMs results in increased ubiquitination of Mtb and reduced bacterial burden (41). Additionally, like Chandra *et al.*, we observed that depletion of *Cyld,* a DUB that counters the activity of the linear ubiquitin assembly complex (LUBAC) (73), resulted in decreased Mtb replication compared to a nontargeting control or scramble. However, in contrast to the Chandra *et al.* study that demonstrated a reduced CFU upon *Usp8* depletion, we observed a modest but reproducible increase in bacterial replication when *Usp8* was knocked down in BV2 cells. The functional divergence regarding *Usp8* between our studies underscores the importance of cellular context (BV2 cells versus BMDM), screen design (luminescent growth assay versus ubiquitination/CFU), and bacterial strain (Mtb Erdman versus H37Rv) when interpreting DUB function. Nevertheless, both studies converge on the therapeutic potential of modulating the host ubiquitin system to combat intracellular pathogens.

Importantly, USP15-IN-1, a small molecule inhibitor, mimics the effects of USP15 deletion in both murine and human macrophages, with enhanced bacterial clearance observed at non-cytotoxic concentrations. The specificity of this effect, absent in USP15-deficient cells, highlights its therapeutic potential. Remarkably, hMDMs were sensitive to much lower concentrations of USP15-IN-1 than murine BV2 cells, despite 95% sequence identity between human and mouse USP15. This observation suggests that modest USP15 inhibition may be sufficient to enhance antimicrobial activity in human macrophages, and that USP15-IN-1 or related compounds may warrant further investigation as host-directed therapeutic agents in combination with antibiotics.

While we were unable to test USP15 function in vivo due to the perinatal lethality of global *Usp15* knockout mice, our results in primary cells strongly suggest physiologic relevance. Future studies using conditional or inducible knockout models of USP15 (74) or administration of USP15 inhibitors with established pharmacokinetics and pharmacodynamics will be essential to determine whether these findings extend to host defense in the context of whole-animal infection.

In conclusion, our data define USP15 as a key negative regulator of bacterial xenophagy in both mouse and human macrophages and establish proof-of-concept that pharmacologic inhibition of USP15 enhances control of Mtb in human macrophages. These findings have implications for understanding the roles of deubiquitinases in innate immunity against intracellular pathogens and support further development of USP15-targeted strategies as a component of host-directed therapy for TB and potentially other infections.

## Materials and Methods

### Bacterial Strains

We used the Mtb Erdman strain for all Mtb experiments. The Mtb mCherry-expressing strain was previously described (22). We received the pLux plasmid (pMV306hsp+LuxG13, BEI plasmid #26161) as a gift from Brian Robertson and Siouxsie Wiles and electroporated it into Mtb Erdman as described (75). We cultured all strains in 7H9 medium supplemented with 0.5% glycerol, 0.05% Tween 80, and 10% Middlebrook OADC (BD Biosciences), as previously described (22). For the USP15-IN-1 growth curve, we added 0 µM, 3.75 µM, 15 µM, or 60 µM of inhibitor to 7H9. We diluted Mtb to an initial OD_600_ of 0.15 and then exposed to USP15-IN-1 for four days.

### Cells and Cell Culture

#### Mouse cells

We cultured the BV2 murine microglial cell line in DMEM (Gibco, 11965-092) supplemented with 10% fetal bovine serum (FBS; Gibco, 16000-044) and 1% HEPES (cytiva, SH30237.01). For the initial screen, because the efficacy of each shRNA was unpredictable, we generated three unique stable BV2 cell lines per gene, each expressing a distinct shRNA sequence (Supplemental Table 1). The same BV2 cell line transduced with a non-targeting shRNA served as the control. Cells were infected with Mtb-pLux at an MOI of 3 to ensure strong luminescence signal. BV2 *Usp15* KO cells were obtained from GEiC Washington University using the gRNA sequence ‘TCTTATAAGCAGTATATGACNGG’. BMDMs were extracted from mouse femurs and tibias and differentiated in DMEM supplemented with 30% L929 cell-conditioned media, 20% FBS, and 1% HEPES (22). After seven days, differentiated BMDMs were harvested for experiments.

#### Human cells

Buffy coats from anonymous donors were obtained from a local blood bank. PBMCs were isolated using SepMate-50 tubes (Stemcell Technology, 85450) following the manufacturer’s protocol. CD14-positive cells were selected using CD14 Microbeads (Miltenyi Biotec, 130-050-201). Adherent monocytes were differentiated in RPMI supplemented with 1% HEPES, 1% sodium pyruvate, 10% heat-inactivated human serum, 10% FBS, and 50 ng/mL GM-CSF (Peprotech, 300-03) for four days. On day five, the media was changed to RPMI with 10% FBS.

##### Mice

Usp15^−/-^ mice on a C57BL/6J background were obtained from Dr. Yue Xiong with permission from Taconic (TF2834). All experiments used Usp15^−/-^ mice and littermate controls from heterozygous crosses. Mice were housed under specific pathogen-free conditions. All studies were approved by the IACUC of UT Southwestern, an AAALAC-accredited institution.

### Lentiviral Transduction for Knockdown in BV2 cells

shRNAs targeting mouse DUBs were obtained from Sigma MISSION (Supplemental Table 1); a non-targeting sequence served as the control. Lentiviruses were generated by transfecting HEK293T cells with lentiviral vectors and packaging plasmids (psPAX2 and pMD2.G). After three days, viral supernatant was collected and filtered through 0.45 µm filters. BV2 cells were exposed to lentiviral supernatant diluted in DMEM and incubated at 37°C with 5% CO_₂_. After infection, the media was replaced with fresh DMEM containing 4 µg/mL puromycin (Fisher Scientific, BP2956-100). Selection was maintained for at least seven days, with media changes every other day.

### Lentiviral Transduction for Knockdown in hMDM

CD14+ cells were differentiated as described above. On day three of differentiation, lentivirus containing USP15-targeting shRNA or non-targeting control was added. Cells were centrifuged for 1 hour at 800 × g. The next day, the media was changed to RPMI with 2 µg/mL puromycin, 1% HEPES, 1% sodium pyruvate, and 10% FBS. Selection was continued for three days. On the day of infection, media was replaced with RPMI supplemented with 1% HEPES, 1% sodium pyruvate, and 10% FBS.

### Macrophage Infections

BV2, BMDMs, and hMDMs were infected with Mtb as previously described (22). Briefly, Mtb cultures were grown to OD_600_ 0.4–0.6, washed three times, and sonicated for three rounds of 7 seconds on and 7 seconds off (total 21 seconds). OD_600_ was remeasured and bacteria diluted to an MOI of 1 for all CFU and immunofluorescence experiments. For Mtb-pLux experiments, an MOI of 3 was used to ensure sufficient luminescence. BV2 cells were seeded at 10 cells per well in white 96-well plates (ThermoFisher Scientific, 165306). Mtb was centrifuged onto cells for 10 minutes at 1500 rpm and incubated for 30 minutes at 37°C with 5% CO_₂_. Plates were read at days 0, 1, 2, 3, and 4. For CFU, BMDMs and hMDMs were seeded at 10 cells/well in 48-well plates (Corning, 3548). For immunofluorescence, 2 × 10 cells were seeded on sterile coverslips in 24-well plates. PIK-III (MCE, HY-12794) was added to inhibit autophagy immediately post-infection. USP15-IN-1 (MCE, HY-148046) was added at 0 µM (DMSO control), 3.75 µM, 15 µM, or 60 µM. Cytotoxicity was measured using the CyQUANT LDH assay (Fisher Scientific, C20300).

### Immunofluorescence

Sixteen hours post-infection with mCherry Mtb, cells were washed twice with PBS and fixed with 4% paraformaldehyde in PBS for 4 hours (26). After fixation, we washed and permeabilized the cells. For LC3, cells were permeabilized with 100% methanol for 5 minutes and blocked with 3% BSA for 30 minutes at room temperature. For K63 and K48 staining, cells were permeabilized with 0.5% saponin for 30 minutes and blocked in 3% BSA with 0.5% saponin at room temperature. Primary antibodies were diluted and incubated at room temperature for 1 hour: anti-LC3 (Sigma, L7543-200UL) at 1:250; anti-K63 (EMD Millipore, 05-1306) and anti-K48 (EMD Millipore, 05-1307) at 1:1000. Secondary antibody (anti-mouse AlexaFluor 488, Invitrogen, A11008) was diluted 1:2000 and incubated for 1 hour at room temperature. Coverslips were mounted with 5 µL ProLong Diamond with DAPI and dried overnight. Images were acquired using a Nikon W1 spinning disk confocal microscope and analyzed using Imaris 10 (Bitplane); image panels were assembled using ImageJ.

### Immunoblots

BMDMs, hMDMs, and BV2 cells were seeded at 10□ cells/well in 6-well plates and either infected at MOI 5 or left uninfected. After 24 hours, cells were washed three times with PBS and lysed in RIPA buffer with protease inhibitors (Roche, 11836170001) for 5 minutes. Lysates were homogenized by pipetting and scraping, then filtered twice through 0.22 µm filters. Protein concentration was determined using a BCA assay (ThermoFisher, 23225). Lysates (20 µg) were separated on 8–20% SDS-PAGE gels (Bio-Rad, 4561096) and transferred to PVDF membranes (Bio-Rad, 1620174) using semi-dry transfer (Bio-Rad, 1704150). Membranes were blocked in 3% BSA in TBST for 1 hour at room temperature. Primary antibodies were used at the following dilutions in 3% BSA: anti-USP15 (Novus Biologicals, H00009958-M01) 1:500, anti-PARKIN (Cell Signaling, 4211) 1:2000, anti-B-ACTIN (Santa Cruz, sc-47778) 1:10000, and anti-LC3B (Novus Biologicals, NB100-2220) 1:250. After three washes, membranes were incubated with HRP-conjugated secondary antibodies (Jackson ImmunoResearch, 115-035-003 or 111-035-003), washed again, and developed using Clarity ECL Substrate (Bio-Rad, 1705060). Blots were imaged using a Bio-Rad ChemiDoc MP system.

### Statistical Analysis

Statistical analyses were performed using GraphPad Prism 9 (version 9.5.0). For CFU and Mtb-pLux experiments involving multiple comparisons, ordinary two-way ANOVA followed by Tukey’s multiple comparisons test was used. For comparisons between two groups, Student’s t-test was used. For co-localization experiments involving more than two groups, ordinary one-way ANOVA was used. All experiments were independently repeated at least three times.

## Authorship contribution statement

KCR: Conceptualization, Formal analysis, Investigation, Writing – original draft, Writing – review and editing, PCC: Investigation, Formal Analysis, Writing – review and editing, BRD: Investigation, Formal Analysis, Writing – review and editing, PKP: Investigation, Formal Analysis, Writing – review and editing, MUS: Conceptualization, Formal analysis, Funding acquisition, Project administration, Supervision, Writing – original draft, Writing – review and editing.

## Declaration of competing interests

All authors declare that they have no competing interests.

## Acknowledgments

The authors would like to thank all members of the Shiloh lab for their support and constructive feedback on the manuscript. We also thank members of the Animal Resources Center at UT Southwestern for providing support with all animal welfare and husbandry. We thank Dr. Yue Xiong, formerly of the University of North Carolina, for the Usp15^−/-^ mice. We would like to thank Dr. Kate Luby-Phelps and Dr. Marcel Mettlen of the Quantitative Light Microscopy Core Facility at UT Southwestern for their assistance with fluorescence microscopy. The spinning disk confocal microscope was funded through the NIH 1S10OD028630 grant to Dr. Kate Luby-Phelps. This work was funded by NIH T32HL098040 (K.C.R.) and 5U19AI142784 (M.U.S.) grants. Michael Shiloh would also like to acknowledge support from the Disease Oriented Clinical Scholar program at UT Southwestern.

## Notes

### Competing Interest Statement

The authors have declared no competing interest.

