## Supplemental Figures and Legends for "Deubiquitinase USP15 restricts autophagy and macrophage immunity to *Mycobacterium tuberculosis*"

### Supplemental Figure 1

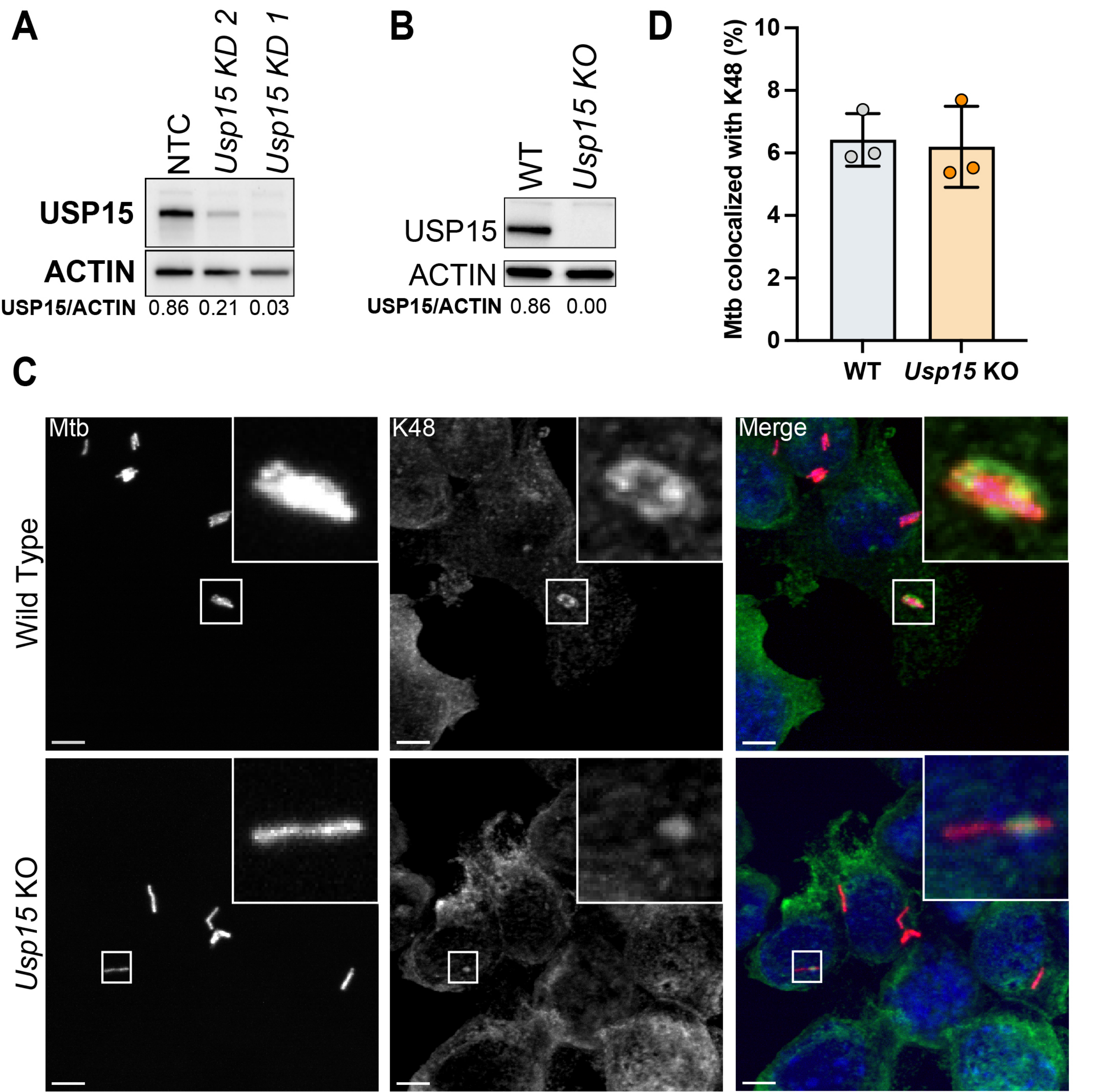

### Supplemental Figure 2

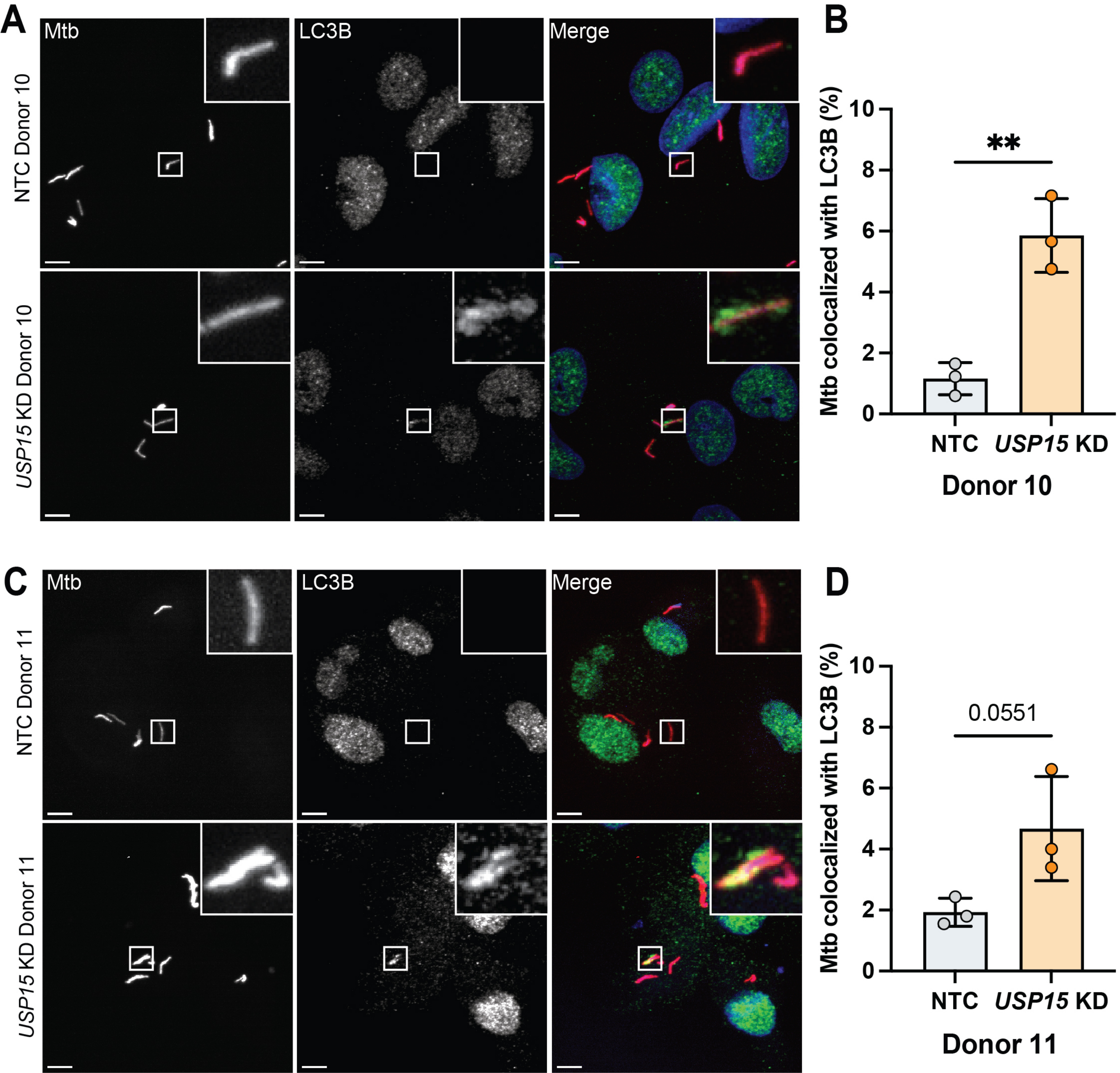

### Supplemental Figure 3

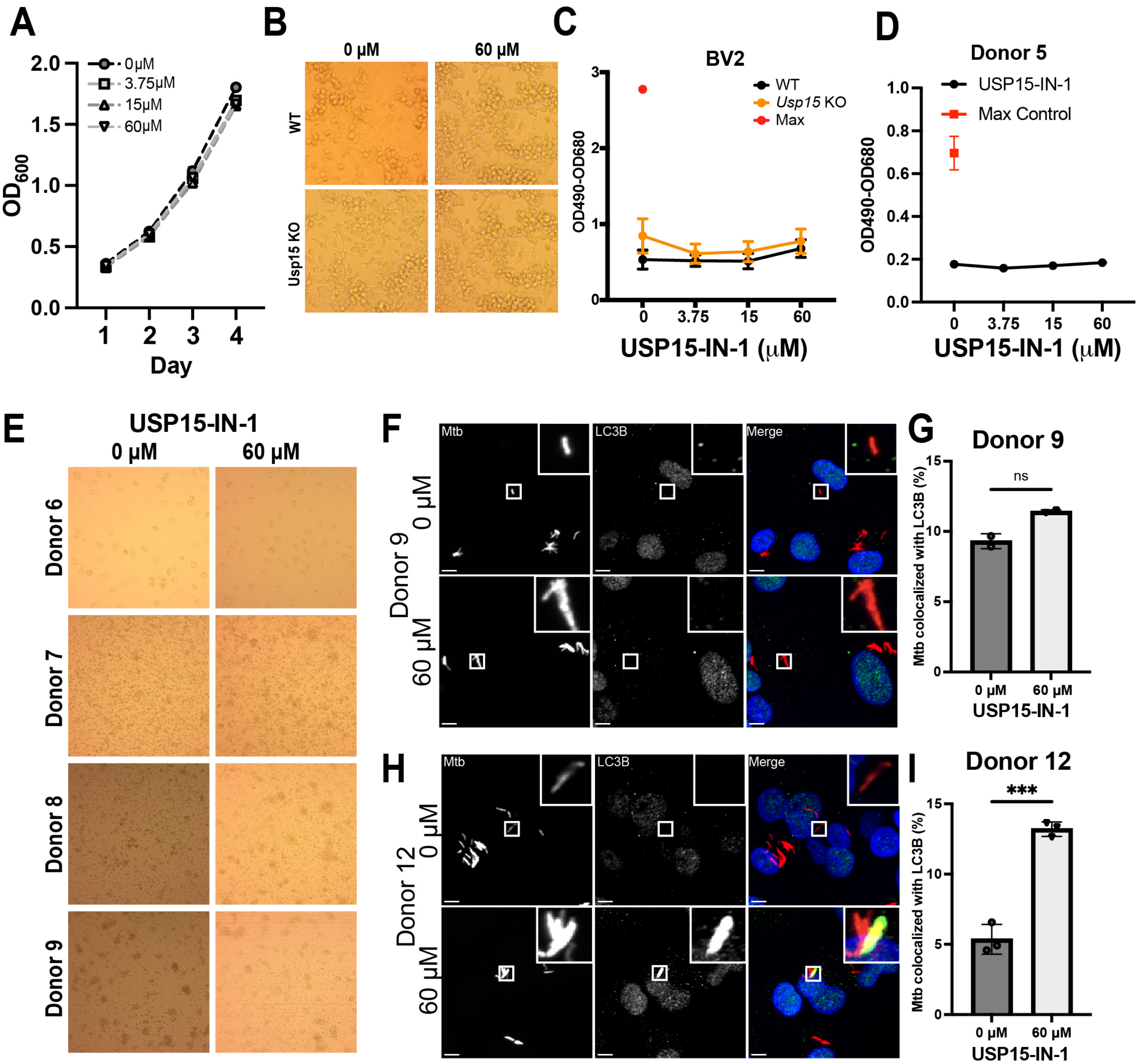

**Supplemental Figure 1.** Validation of USP15 loss and its effect on K48 ubiquitination. A) Western blot of USP15 in BV2 NTC cells and in two of the three BV2 *Usp15* knockdown (KD) cells. B) Western blot of USP15 in WT or *Usp15* KO BV2 cells. C) Representative immunofluorescence images of K48-Ub with mCherry Mtb in WT BV2 cells. Scale bar is 5 µM. D) Quantification of K48 co-localization with mCherry-expressing Mtb in WT or *Usp15* KO BV2 cells. Data shown are representative of at least 3 independent experimental replicates.

**Supplemental Figure 2.** USP15 depletion in hMDMs leads to increased LC3 co-localization with Mtb-associated structures in two donors. A,C) Representative immunofluorescence images of mCherry Mtb (grey or red) and LC3 (grey or green) in hMDMs from A) Donor 10 or C) Donor 11 with NTC or *USP15* KD. Scale bar is 5 µM. B,D) Quantification of LC3 co-localization in hMDMs from B) Donor 10 or D) Donor 11 with NTC or *USP15* KD. For statistical analysis, we used Student's t-test.** $p<0.01$.

**Supplemental Figure 3.** Impact of USP15-IN-1 on BV2 and hMDM viability and LC3 colocalization with Mtb-associated structures. A) Growth curve of Mtb Erdman in 7H9 with different concentrations of USP15-IN-1. B) 10X image of BV2 WT or BV2 *Usp15* KO exposed to DMSO or 60 µM USP15-IN-1 at day 3 after Mtb infection. C,D) LDH assay showing the 4-point dose response of C) BV2 WT and BV2 *Usp15* KO cells or D) hMDMs treated with USP15-IN-1 at day 3 after Mtb infection. E) 10X images from each donor in the RLU experiments with 0 µM (DMSO) or 60 µM USP15-IN-1. F, H) Representative immunofluorescence images of mCherry Mtb (grey or red) and LC3 (grey or green) in hMDMs from F) Donor 8 or H) Donor 12 treated with 0 µM (DMSO control) or 60 µM USP15-IN-1 at 18 hours post-infection. Scale bar is 5 µM. G, I) Quantification of LC3 co-localization in hMDMs from G) donor 8 or I) donor 12. For statistical analysis, we used Student's t-test. *** $p<0.001$. For non-human donor experiments, data shown are representative of 3 independent experimental replicates.
